# Cytoplasmic self-organization established by internal lipid membranes in the interplay with either actin or microtubules

**DOI:** 10.1101/506436

**Authors:** Sindy K. Y. Tang, Malte Renz, Tom Shemesh, Meghan Driscoll, Jennifer Lippincott-Schwartz

## Abstract

Cells harbor an intrinsic organization of their components. Specific protein structures, as the centrosome, have been described master regulators of cell organization. In the absence of these key elements, however, cytoplasmic selforganization has nevertheless been observed. Cytoplasmic self-organization was postulated to arise from the interaction of microtubules with molecular motors on lipid membrane surfaces.

Here, we show that lipid membranes are capable of organizing both major cytoskeletal systems, microtubules and actin, even if one or the other cytoskeletal system is completely paralyzed. A microfluidic droplet system and Xenopus oocyte extracts enabled us to build an artificial cell and study minimal requirements for cellular self-organization. Mathematical modeling reveals the interaction of lipid membranes with any filament system through molecular motors as a universal principle of cytoplasmic self-organization. Both cytoskeletal systems form mechanisms to establish robust 2-dimensional selforganization and self-centering. Pharmacologic inhibition of the cytoskeletal network systems helps dissect specific contributions of each network in the interplay with lipid membranes with regards to 2- and 3-dimensional organization, time and length scale of cytoplasmic organization and the degree of concentration of the centered elements. While microtubules provide 3-dimensional polarity, actin filaments ensure fast and dense compaction and long-range organization.

## Introduction

The mammalian cell displays a typical conserved architecture of its main constituents, cell nucleus, internal membranes, and cytoskeleton. The cell nucleus and adjacent nuclear, and ER / Golgi-associated membranes localize in the cell center with the microtubule cytoskeleton extending outward forming a radial array, while the actin cytoskeleton is predominantly peripherally enriched. Given this fundamental cellular organization, the question arises as to how a cell is able to establish and maintain its architecture. How does a cell define its center and organizes its cellular components? The centrosome, a major microtubule-organizing center in animal cells, has been assumed the critical regulator for cellular organization [ref]. However, cellular organization in the absence of this master regulator has been observed. In surgically cut fish melanophore fragments that lack centrosome, pigment granules aggregate towards the geometrical center of the melanophore fragment upon adrenaline stimulation and a radial array of microtubules forms extending outward. Such cytoplasmic organization occurs in the absence of centrosome, indicating that the centrosome is not the only master of organization in the cytoplasm [1-3]. Apparently, beside the centrosome additional organizing principles need to be considered that have been summarized as principles of cytoplasmic self-organization and self-centering [4, 5]. Critical additional organizing principles appear to be microtubules and internal membranes. It has been proposed that self-organization arises from the “interaction of dynamic microtubules with motors and the surface membrane” [6-9]. Recent studies in intact mammalian cells showed that in fact internal membranes of the Golgi apparatus can function as unconventional microtubule-organizing centers [10, 11] as observed during the reassembly of Golgi mini-stacks after mitotic break down [12-14]. Further organizing principles of self-organization remain to be characterized. Even though cross talk between the actin and the microtubule cytoskeleton that coordinates and facilitates vesicle transport has been reported [15-17], the role of the actin cytoskeleton in the process of cytoplasmic self-organization is yet to be elucidated. Importantly, no study has addressed cytoplasmic self-organization without any pre-existing intrinsic organization.

Here, we show that internal lipid membranes are capable of organizing both cytoskeletal filament systems, microtubules and actin, and establish cytoplasmic selforganization and self-centering. Using microfluidic droplets and mathematical modeling, we demonstrate mechanisms of centering that comprise the interaction of internal lipid membranes with either cytoskeletal filament system. No higher-order regulator is needed. While the concerted interplay of actin and microtubules guarantees robust centering of internal membranes and lipid organelles in the process of cytoplasmic selforganization, each cytoskeletal system contributes unique features; actin mediates long-range compaction, microtubules introduce axial asymmetry.

## Results

### Microfluidics generated droplets for studying cytoplasmic self-organization

With a microfluidics-based droplet system (see methods), we studied the selforganization of cytoplasm derived from *Xenopus laevis* egg extracts in interphase (see Methods). The use of microfluidics allows generation of a large number of droplets with defined dimensions from tens of microns to a few hundreds of microns. These droplets can act as miniaturized compartments for biological processes and thereby provide a platform to study fundamental properties of the cytoplasm. Making droplets that approximate the size of a typical mammalian cell like a fibroblast provides appropriate boundary conditions for the biomechanical activities of actin and microtubules as they assemble and disassemble. The surfactant [18] and oil used here are biocompatible and highly permeable to gases. The generated droplets also allow for high-resolution confocal imaging to view membranes, actin and microtubules in three-dimensions. Storing multiple droplets in a microfluidics device permits observation of multiple samples at the same time.

For generating droplets containing *Xenopus* egg extract, we used two approaches. Gently shaking cell extract in surfactant solution breaks the extract into droplets of various sizes simply by shear (Figure 1A) [19]. Although the size distribution of such generated droplets is wide, this method is straightforward and does not require sophisticated equipment or infrastructure. Alternatively, we used a microfluidic droplet generator [20, 21] to create droplets of uniform size providing high-throughput capabilities [22]. The *Xenopus* egg extract-containing droplets are injected into a microfluidic chamber that has been sealed on a cover slip to allow for confocal imaging. This microfluidic chamber consists of pockets with diameters of 250 μm that are connected by narrow orifices with a width of 50 μm. The chamber height is 40 μm (see Material and Methods for details). At steady state, the majority of the droplets with a diameter of about 250 μm or larger settles inside the pockets to minimize surface energy. Droplets adopt the form of a pancake with a curved lateral surface and flattened top and bottom boundaries. With either approach, we focused on droplets with a diameter of 100 - 200 μm and height ~ 40 μm, as this approximates the typical size of a mammalian cell like a fibroblast.

**Figure 1:**
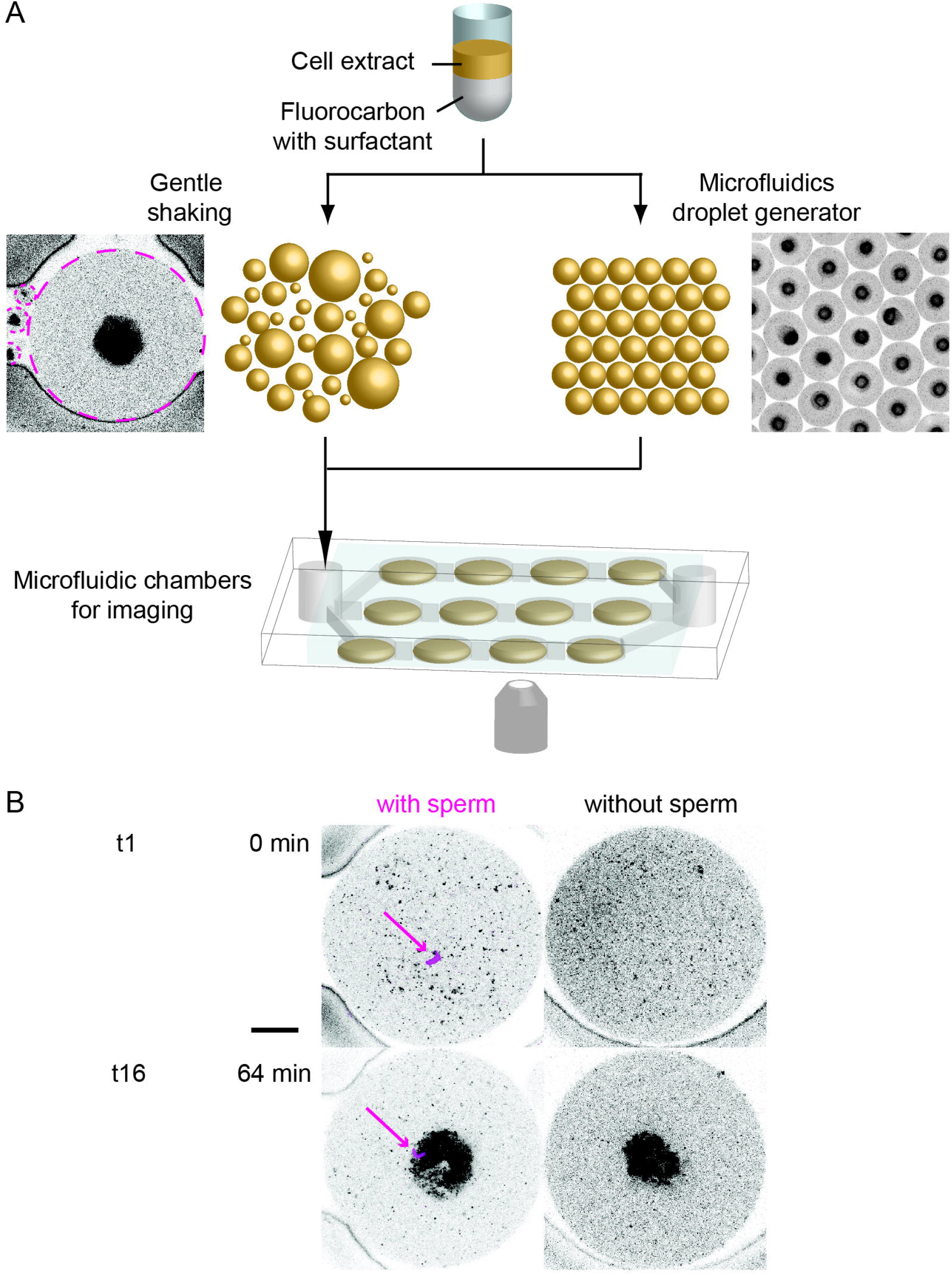
Microfluidics droplet generation, formation of drops containing cytoplasmic *Xenopus* egg extract, and observation of organization and centering of internal membranes in the presence and absence of centrosome (sperm). **Figure 1a:** Scheme outlines different options to generate droplets. Gently shaking cell extract in surfactant solution breaks the extract into droplets of various sizes by shear. Although the size distribution of such generated droplets is wide, this method is straightforward and does not require extra equipment. Alternatively, a microfluidic droplet generator can be used that creates droplets of highly defined and uniform size. Microfluidics allows generation of up to 10^3^ droplets or vesicles per second. **Figure 1b:** The final organization of internal membranes in the center of a droplet is identical in the presence and absence of centrosome (sperm) as assessed by confocal imaging Scale bar 50 μm.

Freshly extracted cytoplasm from eggs in interphase shows no preformed organization; microtubule and actin cytoskeletal fibers are disrupted and internal membranes fragmented to small vesicles. Within one hour, however, the cytoskeletal fibers polymerize and the vesicles center to the middle of the droplet (Fig. 1A). Thus, our approach permits visualization of the initial nucleation and polymerization of microtubules and actin filaments, changes in membrane distribution, and the gradual emergence of cytoplasmic self-organization in interphase over time.

### Centering of cytoplasmic components does not require centrosome

We introduced sperms into the egg extract by pre-mixing the solutions of known concentrations and volumes prior to droplet generation. The incorporation of sperm and thereby the centrosome as microtubule-organizing center into egg extract was, thus, a stochastic process; with many droplets comprising sperm/centrosome and some without. We noticed, however, that after approximately 30 min, all droplets exhibited centering of internal membranes surrounded by a ring of microtubules, regardless whether they contained sperm/centrosome or not (Figure 1 A). A side-by-side comparison of a droplet containing sperm/centrosome and a droplet without centrosome confirmed that the final organizations of DiI labeled internal membranes were the same in both cases as assessed by confocal imaging (Figure 1B). We concluded that for cytoplasmic organization and centering in interphase, the centrosome as microtubule-organizing center is not critical at the level of our observation. We therefore set out to study the fundamental properties of cytoplasmic self-organization in the absence of centrosome in detail using labeled tubulin, utrophin-GFP-labeled actin, and DiI-labeled internal membranes (see Material and Methods).

### Temporal and spatial pattern of cytoplasmic self-organization in microfluidic droplets

Immediately after the formation of microdroplets containing cytoplasmic extract, internal membranes were randomly distributed throughout the droplet (Figure 2). We detected few polymerized microtubules and no actin filaments. Initially, microtubules appeared to nucleate or polymerize randomly in all directions throughout the droplet. Slowly, microtubule filaments merged to form micro-asters. Concurrently, internal membranes started to accumulate at the center of the droplet. As the membranes became more clustered towards the center, filamentous and dimeric tubulin cleared out from the center and became radially distributed forming a dense ring of polymerized filaments around the centered membranes radiating outwards. The spatial relationship between microtubules and lipid organelles was consistent with these organelles serving as a surface for microtubule polymerization and mutual movement of lipid organelles and microtubules alongside each other, consistent with previous experiments (Rodionov, kimura).

**Figure 2:**
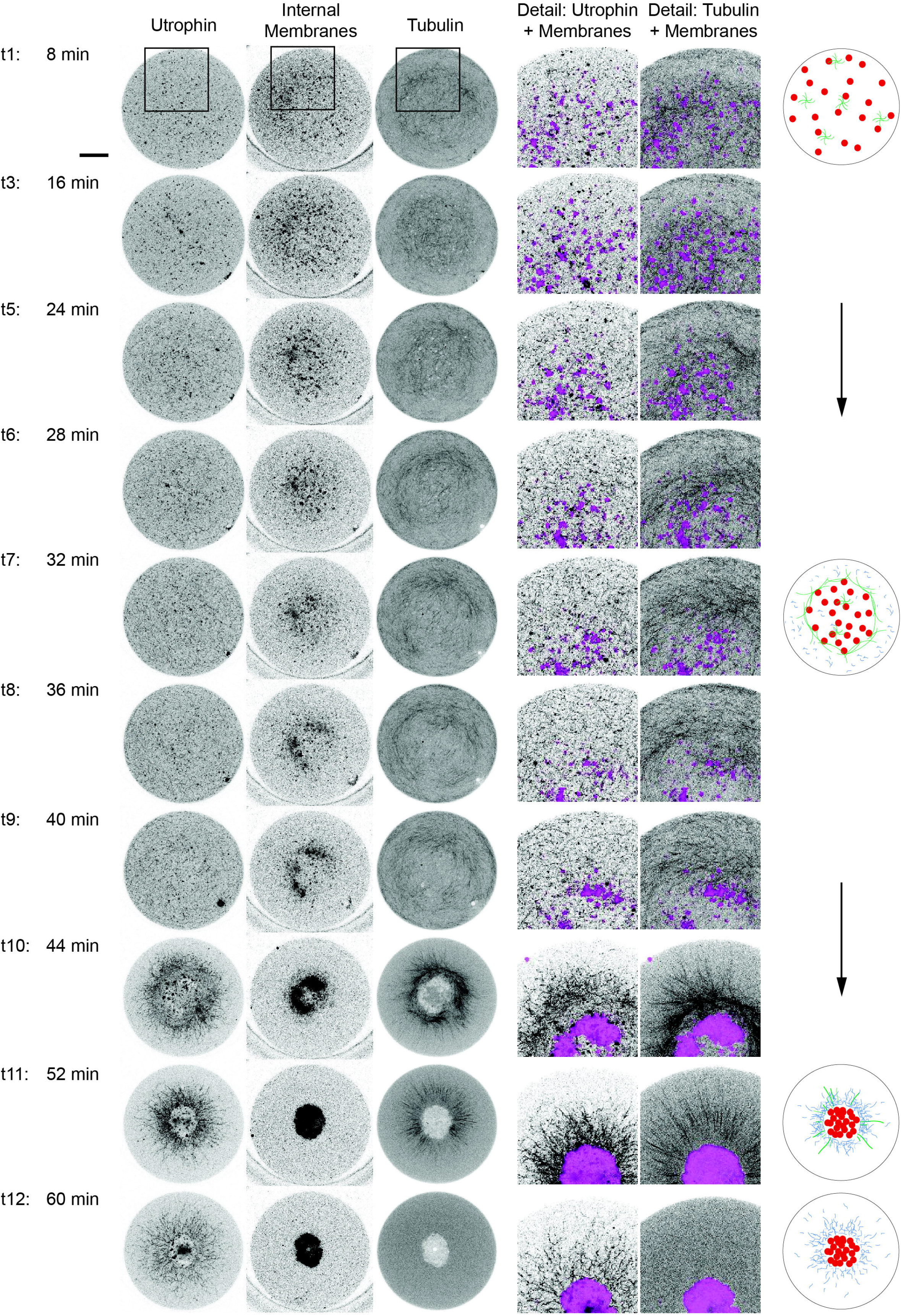
Time course of cytoplasmic self-organization in microfluidic generated droplets. Initially evenly dispersed internal membranes become gradually centered over time. Initially, microtubules nucleate randomly throughout the droplet. Slowly, microtubule filaments grow and merge to micro-asters. Concurrently, internal membranes accumulate at the center of the droplet. As membranes become more clustered towards the center, tubulin clears from the center and becomes radially distributed forming a dense ring around the centered membranes radiating outwards. After 16 – 20 min, polymerized actin can be visualized. It polymerizes predominantly in the outer droplet region that has already been reduced in lipid organelles. In contrast to the microtubule system, the actin filaments are short and highly branched forming a cortical meshwork. This meshwork contracts at about 40 min and induces microtubule filaments irreversible shortening, known as catastrophe[23][23][23][22][21][20][19]. In this final stage, internal membranes of lipid organelles are compacted in the droplet center. Purified GFP-utrophin is used for labeling actin (left panel), DiI for labeling Golgi and ER derived internal membranes (middle panel), and purified alexa647-tubulin for labeling microtubules (right panel). In the schematic, microtubules are depicted in green, actin in blue, and internal membranes in red. Scale bar 50 μm.

Notably, after 15 – 20 min, short polymerized actin filaments could be visualized which showed no preferred directionality but were rather short and highly branched. Over time, the polymerized actin filaments seemed to dominate in the outer region of a droplet with a reduced number of lipid organelles. This circular band of high-density actin appeared to collapse inward to the center of the droplet at about 40 minutes. Concurrently, an irreversible shortening of the microtubule polymers was detected that reminded of microtubule catastrophe [23]. Eventually, no filamentous microtubules were visible. In this final stage, internal membranes of lipid organelles were highly compacted in the center of the droplet with dimeric tubulin as well as actin largely excluded from this region.

Both cytoskeletal filament systems, i.e. microtubules and actin, are apparently contributing to the observed cytoplasmic self-organization in the absence of centrosome. Both cytoskeletal filament systems drive the initially randomly distributed internal membranes to the center of the microfluidic droplet. In the following, we generate a general model comprising cytoskeletal filaments, motors and internal lipid membranes and the interaction between the three components. We hypothesized an overarching basic principle of both cytoskeletal systems in the absence of a master regulator and test if such interaction would be sufficient to simulate our observations.

### General model of internal membrane and cytoskeletal filaments interaction predicts cytoplasmic self-organization

In our experimental set-up, the frog egg extracts directly after encapsulation into microfluidic droplets show no preformed cytoskeletal filaments or only short filaments with no orientation and there is no microtubule-organizing center such as centrosome. Thus initially, the frog egg extract is a rather homogeneous mixture of internal membranes, mono- and dimeric cytoskeletal components and only short cytoskeletal filaments. In our model, the basic assumption is that polymerizing cytoskeletal elements can transiently bind via molecular motors to internal membranes fragments. Due to the polar nature of cytoskeletal filaments, molecular motors advance along the filament towards a preferred polar tip; resulting in an attractive force between the membrane fragments and the target tips of the filaments. Together, filaments, and internal membranes form short-lived contractile elements. Each component of the system (membranes, filaments) experiences pulling forces from the complementary component (filaments, membranes) immediately surrounding it, while the range of this attractive force is determined by the length of the cytoskeletal filaments. For any membrane fragment in the system, the magnitude of the pulling force applied in each direction is determined by the concentration and orientation of filaments in the surrounding region. The combined pulling forces experienced in a specific region result in a flow in the direction of the net force. Similarly, the average force exerted on any filament is determined by its orientation, as well as the distribution of membrane fragments in its vicinity. The motion of cytoskeletal / membrane elements is therefore determined by a combination of (i) free diffusion and (ii) contractility induced flow.

From these considerations, we have constructed a mathematical model for the time evolution of the concentrations of the membrane fragments, *c_m_*, and cytoskeletal filaments, *c_f_* (Figure 3; for a detailed derivation, see Supplementary Material). Our model takes into account the different diffusion and drag properties of the membranes and the filaments, as well as the mean length of the filament, *L_f_*, which is assumed to grow over time due to filament polymerization.

**Figure 3:**
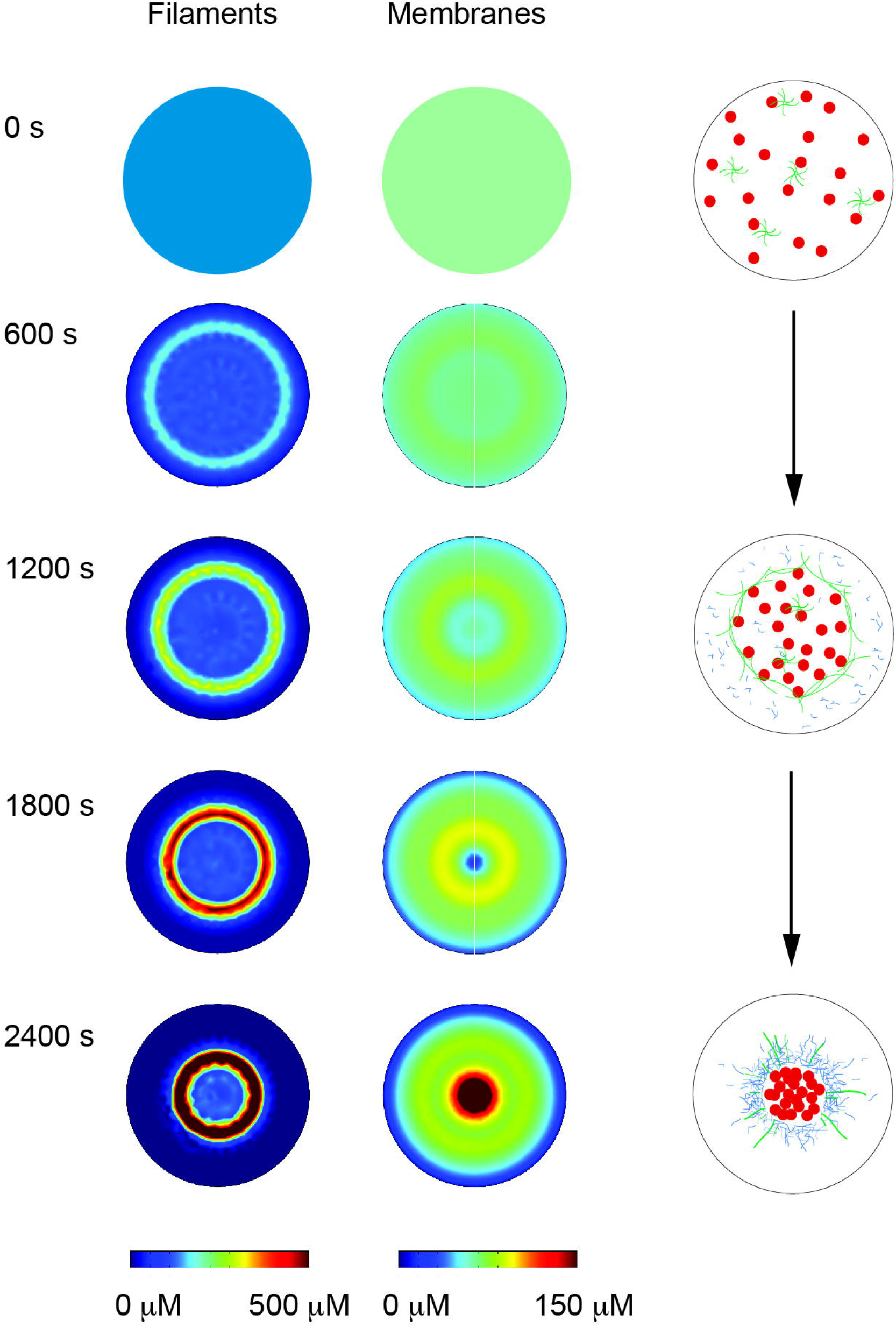
Model simulation of cytoplasmic self-organization based on internal membranes, molecular motors, and cytoskeletal filaments. The spatial distributions of cytoskeletal filaments (bottom) and membrane fragments (top) at different time points are shown. The initial conditions at time 0 s are of a homogeneous distribution of system components, contained in a cylindrical droplet with a diameter of 200 μm. As filaments elongate due to polymerization, they begin to interact with membrane bound motors, resulting in a pairwise attractive force between the membrane fragments and the target filament tips. Initially, the net attractive force is zero throughout the system, except for a region close to the droplet boundary, where a small centripetal force develops. As the system evolves in time, cytoskeletal filaments condense into a ringshaped region, while membrane fragments form a band with a smaller radius. Ultimately, the inner membrane ring coalesces into a circular core at the droplet center, surrounded by membrane fragments. Initial concentrations are taken as 100 μM, with local concentration indicated by color-bars.

For an initial condition of a homogenous distribution of both components, *c_m_* and *cŗ*, then the contractile force is uniform everywhere - except in a region near the droplet boundary. Cytoskeletal filaments and internal membranes close to the boundary experience centripetal pulling forces without a balancing force pulling them outwards, since there are no contractile elements beyond the droplet boundary. Therefore, there is a net flow towards the center, resulting in a peripheral band with elevated concentrations. Importantly, a finite interaction range is required for the development of a force imbalance, and hence the initial band formation is only expected once the filaments reach a threshold length, *L_0_*. Steric interactions between the long cytoskeletal filaments and the droplet edges result in an initial depletion of filaments at the periphery of the droplet, promoting a greater force imbalance and centripetal flow. This steric repulsion and resulting force imbalance become more pronounced for droplet edges with a higher curvature. Since the droplet top and bottom edges are essentially flat, we model the planar flow in the system, neglecting a vertical component.

At later time steps, the coupling between the internal membranes and cytoskeletal filaments results in a contraction of the band, and a simultaneous increase in the concentration within it. As the filaments polymerize and grow in length, the range of the attraction between the membranes and the filaments increases, and consequently the centripetal force grows and the contraction of the ring is accelerated. Our results indicate that if the viscous drag acting on the membrane fragments is greater than the drag on the filaments, then the filament band becomes peripheral to the membrane, ultimately resulting in a membrane core surrounded by cytoskeletal filaments.

Our model demonstrates that simple motor-induced interaction between internal membranes and cytoskeletal elements results in spontaneous formation of distinct morphology and in a global coordinated motion - without the requirement for higher-order coordinating components, such as the centrosome.

While the proposed mechanism for centralization of cytoskeletal elements is, fundamentally, the same for microtubules and actin filaments, differences in the mechanical properties, as well as the different molecular motors associated with each type of filament, can affect the kinetics of the centering motion and the final geometry of the system. Further, our model does not take into account possible interactions between the cytoskeletal sub-systems. The general prediction of our universal model is that internal lipid membranes are capable to organize each individual filamentous system, resulting in the cytoplasmic self-organization we observed in microfluidic droplets.

### Microtubule mediated self-centering lacks final compaction

We hypothesized that paralyzing the actin network should lead to lipid membrane and microtubule based self-organization and centering according to our model. In contrast to the classical experiments in fish melanocytes, in the system presented here no preformed microtubule filaments and no external ligand, i.e. adrenalin, stimulation is needed. Upon addition of latrunculin A, we found microtubules nucleated and polymerized throughout the droplet just as in the initial states of unperturbed cytoplasmic self-organization (Figure 4). However, the microtubule filaments here appeared to be thicker and more bundled than in the unperturbed sequence. As the membranes became clustered towards the center, filamentous and dimeric tubulin cleared out from the center and became radially distributed forming a dense ring of polymerized filaments around the centered membranes radiating outwards. As in the unperturbed sequence, the spatial relationship between internal membranes and microtubules was consistent with organelles serving as a surface for polymerization of microtubules and mutual movement of lipids organelle and microtubules alongside each other results in self-organization and centering. In contrast to the unperturbed state, there was no final compaction and catastrophe of the radial microtubule array. Instead, outward extending filamentous microtubules remained stable for more than 100 minutes. The persistence of highly bundled microtubules over a long period of time is consistent with the idea that the actin meshwork plays a role in providing factors for destabilization and turn-over of microtubules. Similar results have been achieved with cytochalasin D, supporting the notion that the microtubule cytoskeleton is sufficient in centering of lipid organelles and internal membranes but lacks the final tight compaction of internal membranes.

**Figure 4:**
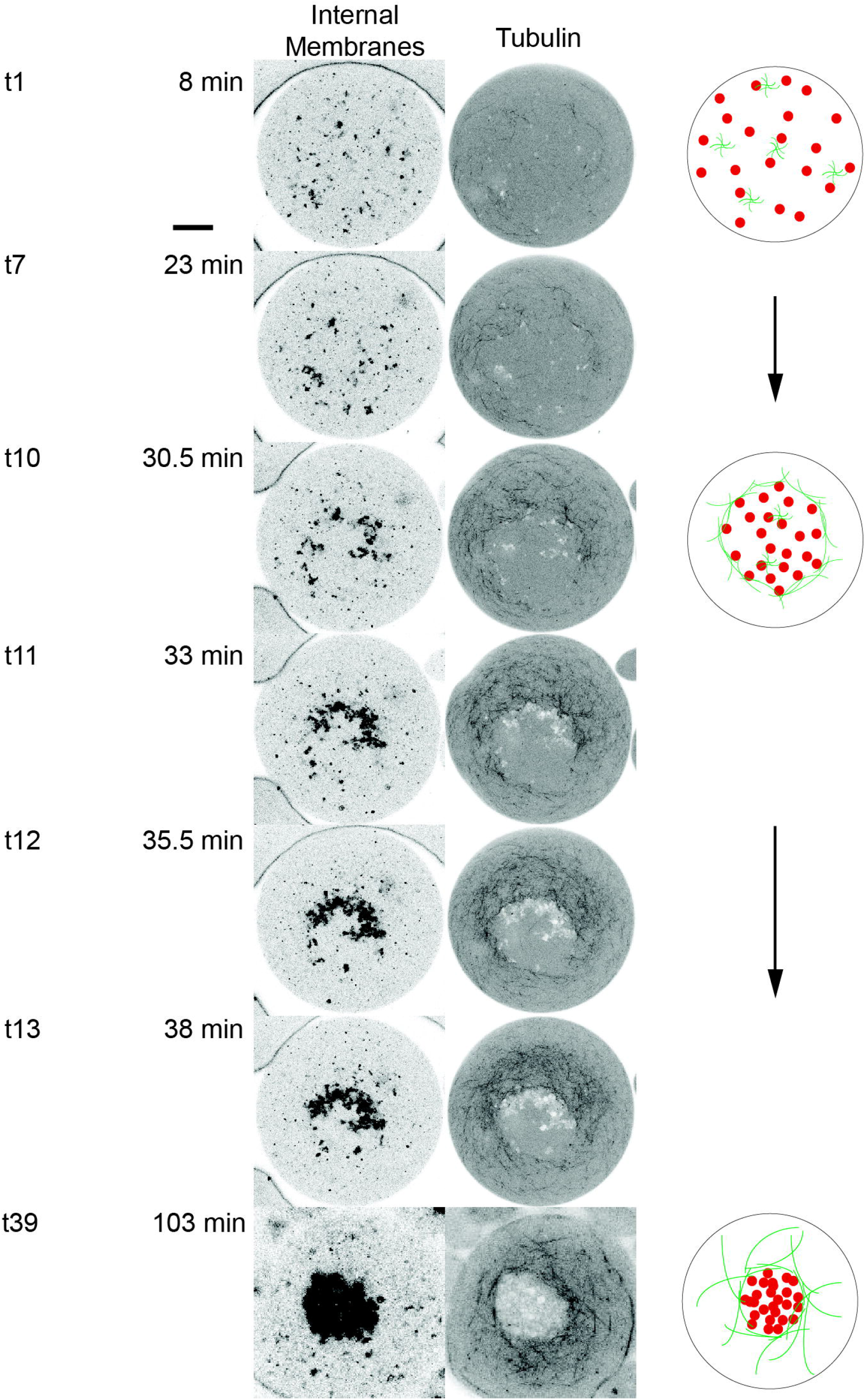
Self-organization of cytoplasm in the presence of latrunculin A, a G-actin sequestering agent. As in the unperturbed sequence, the spatial relationship between microtubules and lipid organelles is consistent with organelles serving as a surface for polymerization of microtubules and mutual movement of lipids organelle and microtubules alongside each other results in self-organization and centering. However, there is no final compaction and catastrophe of the radial microtubule array. Instead, the microtubules remain stable for more 100 minutes. DiI for labeling Golgi and ER derived internal membranes (left panel), and purified alexa647-tubulin (right panel). Scheme outlines self-centering of internal membranes over time; microtubules are outlined in green, and internal membranes in red. Scale bar 50 μm.

### Internal membranes interacting with actin cytoskeleton sufficient for self-centering

Our general model of cytoplasmic self-organization predicts that internal lipid membranes can form contractile elements with cytoskeletal filaments via molecular motors and are capable of cytoplasmic self-organization and centering. Our simulations and our observation that a dominant actin filament meshwork forms during the unperturbed cytoplasmic self-organization led us to hypothesize that actin mediated contraction may in fact be sufficient for self-centering of cytoplasmic components with no microtubule filaments necessary. To test this hypothesis, we added nocodazole to the Xenopus egg extracts to prevent microtubule polymerization and the formation of a radial microtubule array. Under these conditions, we noticed on the homogeneous background of labeled dimeric tubulin polymerization of short actin filaments that appeared to be more pronounced than in the presence of filamentous microtubules (Figure 5). Initially, the emerging short actin filaments showed no preferred orientation and were distributed randomly throughout the droplet. After 28 min, a peripheral enrichment of actin filaments surrounding the lipid organelles was clearly visible. In fact, the lipid organelles appeared to be lined up along the actin meshwork. The emergent actin meshwork grew faster than in the presence of filamentous microtubules and contracted earlier (at about 32 min vs. 45 min in the presence of filamentous microtubules). The resulting centering of lipid organelles took place more rapidly than in the presence of filamentous microtubules (5 min vs. 20 min). In this process, lipid organelles are transported in a short period of time over about 100 μm from the droplet periphery to its center. The absence of polymerized microtubules as balancing resistance may allow the actin meshwork to contract earlier and more rapidly. Although the described actin-based self-organization was different from the centering process observed in the presence of polymerized microtubules, final centering was achieved even in the absence of polymerized microtubules. Of note, when both actin and microtubule polymerization was inhibited by adding latrunculin and nocodazole simultaneously, no centering of lipid membranes was observed (Suppl. Fig. 1)

**Figure 5:**
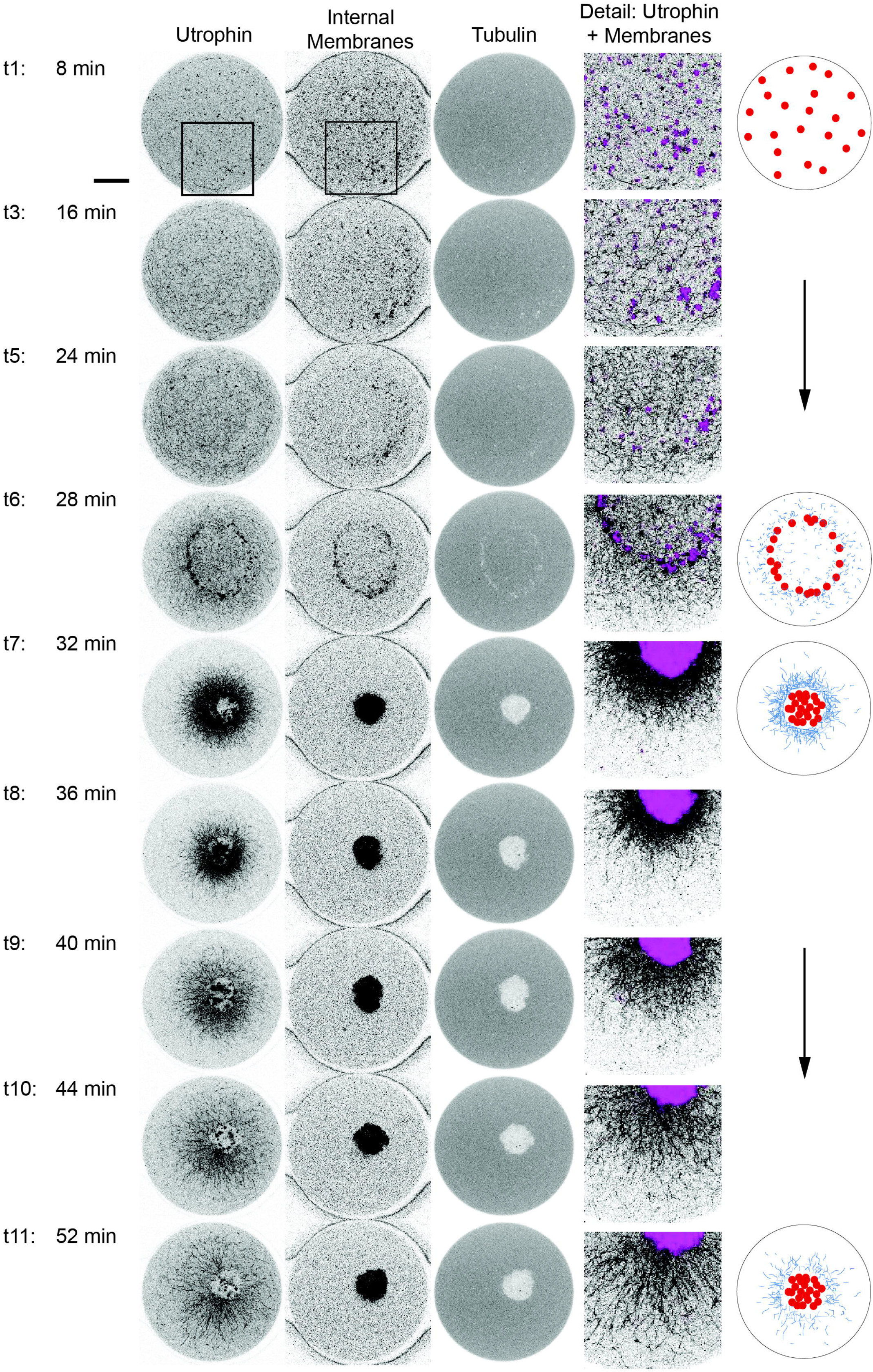
Self-organization of cytoplasm in the presence of nocodazole, a microtubule depolymerizing agent. Only the homogeneous background of labeled dimeric tubulin was detectable. Actin network formation starts again in the outer cortical droplet region, grows faster, and contracts earlier, already after 25 min. Purified GFP-utrophin for actin labeling (left panel), DiI for labeling Golgi and ER derived internal membranes (middle panel), and purified alexa647-tubulin (right panel). In the schematic, microtubules are depicted in green, actin in blue, and internal membranes in red. Scale bar 50 μm.

### The interplay of actin and microtubule networks guarantees robust geometrical centering in 2 dimensions

Having determined that internal lipid membranes can act as organizational principles of the actin and microtubule cytoskeleton in cytoplasmic self-organization, we further analyzed the spatial robustness of cytoplasmic self-centering in two dimensions. In order to test cytoplasmic self-organization in large and asymmetrically shaped droplets, we generated droplets that span more than one pocket in the microfluidic device used and were, thereby, shaped like a dumbbell. As shown in Suppl. Fig. 2, in such a dumbbell-shaped droplet, lipid organelles became initially aggregated in the ‘dumbbell weights’ slightly off-center and closer to the narrow orifice connecting two pockets of the microfluidic device. These aggregates of lipid organelles were surrounded by a radial microtubule array just as in the pancake-shaped droplets. Apparently, the initial microtubule-based self-organization does not result in perfect self-centering. As the initiation of the centering motion is attributed to a force imbalance arising near the droplet edges, the centripetal flow in the region close to the neck of the dumbbell droplet is diminished. This radial asymmetry in the flow of cytoskeletal elements and membrane fragments results in an off-center locus of compaction in the circular regions of the droplets.

Importantly, the actin network on the other hand appears to mediate the final symmetric centering in the middle of the connecting channel. This indicated that in larger droplets of 500 μm and more while the microtubules may not be long enough to reach the droplet boundary, the actin-based centering is capable of mediating such a long-range collective cargo transport. Apparently, the force of actin filament system and its motors are strong enough to merge the two lipid organelle aggregates. Thus, only the interplay of the two major components of cytoplasmic self-organization ensures that lipid membranes and organelles robustly find the geometrical centroid in two dimensions even in large droplets with an asymmetrically shaped boundary.

### Three-dimensional cytoplasmic self-organization

So far, we had only analyzed the spatial self-organization of cytoplasm in the lateral *x*- and *y*-dimensions. Regarding self-organization in the z-dimension, our general model would predict an assembly of internal lipid membranes forming a cylinder spanning the entire depth of the microfluidic droplet. This is simply based on the difference in dimensionality of the boundaries; while the curved membrane constitutes a 2-D system the flat top and bottom membrane form a 1-D system.

We used three-dimensional reconstruction of confocal z-stacks recorded over the entire depth of the droplet to visualize the vertical positioning of cytoplasmic components. As shown in Figure 8, in the unperturbed state, microtubules nucleated and polymerized strongly at the bottom of the droplet and, over time, formed a cage surrounding the internal membranes that was closed at its bottom and open at the top. At this stage, internal membranes of lipid organelles were excluded from the bottom of the droplet. Such clear axial asymmetry was not visualized in case of the actin filament system. Thus, in contrast to the symmetrical centering in the lateral dimensions, a significant asymmetry along the axial axis was noted which is based on our model assumptions an unexpected observation. Considering the surrounding material, we realized that every surrounding surface was made of PDMS except for the cover glass at the bottom. Therefore, we hypothesized the noted axial asymmetry in microtubule-based organization could be due to differences in stiffness of the boundary, which, in turn, caused differential dynamic instability and resulted in an asymmetric radial microtubule array. Alternatively, differences in specific density of microtubules and lipid membranes may explain the axial asymmetry. In order to dissect effects of gravity and differential boundary stiffness, we turned the microfluidic chambers bottom-up and visualized self-organization after 25 min as well as replaced the cover glass by PDMS. In both scenarios, the microtubule cage was closed at the bottom irrespective of the material it was facing (data not shown). Even though boundary stiffness may be a contributing factor, gravity seems to be the more decisive determinant of asymmetrical axial self-organization. Paralyzing the actin filament system does not alter the axial asymmetric assembly of the microtubules. In the presence of latrunculin the polarized microtubule cage remained stable for more than 100 minutes.

In contrast to the axial asymmetry of microtubule formation and internal membranes, actin network formation is symmetrical along the z-axis. In fact, actin seems to predominantly work in *x*- and *y*-dimension and does not centered lipid organelles in z-dimension just as predicted by our model assumptions. When the contracting actin meshwork lead the radial microtubule array into catastrophe, the internal membranes of lipid organelles spread in a narrow cone over the entire depth of a droplet. In the presence of nocodazole, internal membranes were centered by the actin meshwork only in the lateral dimensions and spread in that narrow cone over the entire depth of a droplet.

## Discussion

Cellular organization has long been studied. Besides master regulators of cytoplasmic organization such as centrosome or kinetochore, additional principles have been described and postulated principles of cytoplasmic self-organization. Here, we show cytoplasmic self-organization of interphase extracts from *Xenopus* eggs in the confined compartments of microfluidics generated droplets. These cytoplasmic extracts do initially not show any organization, however robustly self-organize and center internal membranes of lipid organelles.

For this process, no higher-order regulator is needed. Instead, internal lipid membranes appear to be the major player and capable of organizing both the microtubule and actin cytoskeleton. Internal lipid membranes appear to provide nucleating factors and molecular motors and form hubs for microtubule and actin filament polymerization. Internal membranes and cytoskeletal filaments constitute complementary moieties of contractile elements. Thus, membrane fragments and cytoskeletal filaments undergo both free diffusion and, by interaction with their complementary contractile elements, contractility-induced flow. This net motion results in self-centering of lipid membranes, regardless of the cytoskeletal system involved. In the lateral x-y-dimension actin and microtubule cytoskeleton work together in centering internal membranes and make cytoplasmic self-organization a highly robust and conserved process.

In many aspects, however, the two filament systems, microtubules and actin, appear to compete for the internal lipid membranes. Both bind to the membranes, recruit nucleation factors and molecular motors from them and move alongside the membranes. The stirring of evolving microtubules seems to disrupt the emerging actin network and delay its lateral flow, which becomes apparent when microtubules are paralyzed and not functional. Based on their properties each filament system contributes specific features to cytoplasmic self-organization. Microtubules and actin use different molecular motors and exhibit distinct filament length and structure; they form either rather long and straight tracks or a dense dendritic meshwork. While both cytoskeletal filament systems center along the droplet axis, microtubules introduce asymmetry in the axial assembly of internal lipid membranes. Microtubules form a cage around the internal membranes, which is closed at the bottom (i.e. microtubules form at the bottom of the microfluidic droplet and push the internal membranes up). This axial asymmetry is lost as soon as the microtubule filaments disappear and the actin filaments dominate. The exact mechanism of this observed axial asymmetry is not clear, but may be related to differences in the specific density of microtubules and internal membranes and the influence of gravity on the system. Actin-based internal membrane transport, on the other hand, appears to be capable of covering greater distances and thereby guarantees robust axial compaction even in large, irregularly shaped, microfluidic droplets. Thus, actin mediates long-range transport of dispersed cargo. Lenart et al. showed a similar actin-based mechanism pulling segregated chromosomes of starfish oocytes inward during meiosis [24]. When tightly anchored to the plasma membrane, actin is also able to facilitate the reverse process, i.e. outward transport of vesicles, as described by Schuh et al. [16]. Alternative possible mechanisms for actin-mediated self-centering include acto-myosin active-gel theory [25] and self-centering due to interactions between two types of high-order actin structures: radial and transverse filaments [26]. However, such mechanisms require the establishment of large-scale actin structures prior to centralization, which were absent from our experimental system. Besides the wider distances of actin-based transport, the degree of actin-driven membrane compaction is higher than obtained by microtubules. The observed tight actin-driven compaction reminds of the phenomenon of phase separation with all lipid components densely packed in the center surrounded by hydrophilic cytoplasmic components. In fact, the dense packing and sequestration of microtubule nucleating factors in the center prohibiting new nucleation events in the rest of the microfluidic droplet may cause the eventual total loss of filamentous microtubules.

The described mechanisms of self-organization may play a role not only in defining the cellular architecture or reestablishing it after mitotic break down, but also in the formation of specialized cytoplasmic sub-compartments as for example the immunological synapses. Thus, the presented microfluidics approach of cytoplasmic self-organization of *Xenopus* egg extracts in confined compartments helps address and elucidate the fundamental principles of cellular architecture.

## Acknowledgements

We would like to thankxe the organizers of the Physiology Course Woods Hole for providing resources, Drs. Christine Field and Ani Nguyen for their help preparing the frog egg extracts, Dr. Tim Mitchison for labeled tubulin and GFP-utrophin as well as Drs. John Hammer and Xufeng Wu for critical discussions.

## Material and Methods

### Xenopus egg extract

Interphase extracts were prepared as previously described except that cytochalasin D was not used [27]. CaCl was added to end concentration of 0.4 mM to mimic fertilization. For labeling microtubuli, actin, and DNA purified Alexa-647 or Alexa-488 tubulin (1 mg/ml), GFP-utrophin (? mg/ml, GFP fused to an artificial dimer of actin-binding domain of Utrophin [28, 29]), and Hoechst were used, respectively. For labeling internal membranes (including membranes of the endoplasmatic reticulum and the Golgi apparatus), DiI was added and incubated at room temperature for 15 min. Then, cytoplasmic extracts were placed on ice, incorporated into microfluidic droplets and imaged immediately. Time points in Figures 1 - 6 are indicated as measured after shifting cytoplasmic egg extracts from ice to room temperature for confocal imaging. Nocodazole (Sigma Aldrich) and latrunculin A (Sigma Aldrich) were added to the extracts in end concentrations of 16 μM, and 150 nM, respectively.

**Figure 6:**
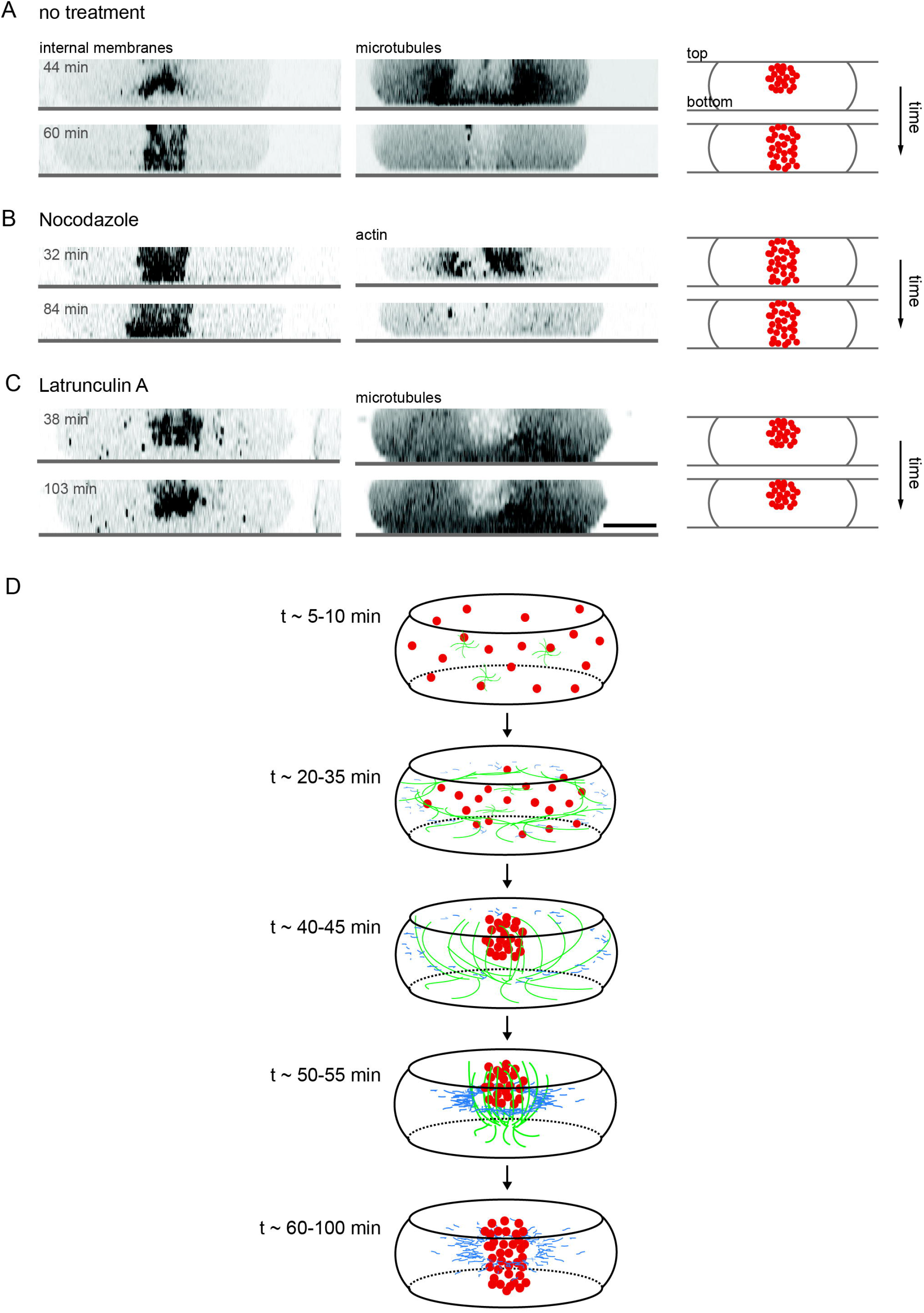
Axial self-organization of cytoplasm without pharmacological perturbation (a) and in the presence of nocodazole (b), and latrunculin A (c), respectively. Internal membranes of lipid organelles are excluded at this stage from the bottom of the droplet. DiI for labeling Golgi and ER derived internal membranes. Scale bar 50 μm. **Figure 6a:** Microtubules nucleate and polymerize more strongly at the bottom of the droplet. They form a cage around internal membranes which is closed at the bottom and open at the top. When the contracting actin meshwork triggers microtubule catastrophe, the internal membranes of lipid organelles spread in a narrow cone over the entire depth of a droplet. **Figure 6b:** In the presence of nocodazole, internal membranes are centered by the actin meshwork only in the lateral dimensions and spread in that narrow cone over the entire depth of a droplet. **Figure 6c:** In the presence of latrunculin, the polarized microtubule cage remains stable for more than 100 minutes. **Figure 6d:** In the scheme of 3-dimensional self-organization of cytoplasm in microfluidic generated droplets, microtubules are outlined in green, actin in blue, and internal membranes in red.

### Microfluidic droplet generation

We fabricated microfluidic droplet generators and imaging chambers in poly(dimethylsiloxane) (PDMS) using standard soft lithography [20, 30]. We used HFE7500 (3M), a fluorocarbon, as the carrier fluid. The surfactant was biocompatible [18] and was necessary to stabilize the drops against coalescence. To generate monodisperse drops of extracts, we used a microfluidic flow-focusing droplet generator [20, 30]. We loaded the extract (pre-mixed with sperms or other labels) into a 1 mL syringe. We used a syringe pump to inject the extract into the microfluidic device. The drops were collected in an Eppendorf tube, and subsequently injected into an imaging chamber using pipettes. Alternatively, drops (with uncontrolled size distribution) were generated by shaking an Eppendorf tube containing equal volumes of extract and carrier fluid containing the surfactant.

### Confocal microscopy

Xenopus egg extract imaging was performed on a Zeiss LSM 700 confocal laser scanning microscope equipped with a 40x Plan Neofluar NA 1.3 oil immersion objective. The 488-nm line of an argon ion laser (Lasos), 555-nm and 639-nm HeNe diode lasers (Lasos) were used to excite GFP/Alexa488, DiI, and Alexa-647, respectively. Dichroic mirror on the LSM 700 was 488/555/639. Fluorescence from GFP and Alexa-488 was collected through a 505 - 530 nm band-pass, fluorescence from DiI through a 585 - 615-nm band-pass filter, and fluorescence from Alexa-647 through a 640-nm long-pass filter. Zen software was used for three-dimensional reconstruction of recorded z-stacks.

### Mathematical modeling

We model the distribution of the system’s components: concentration of cytoskeletal filaments, *c_f_*, and concentration of membrane fragments *c_m_*. The time evolution of the concentrations is described by the standard Smoluchowski equations:

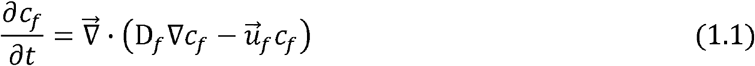

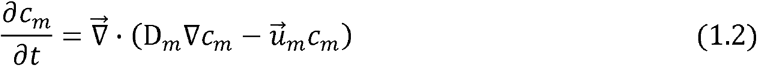

Where D_*f*_ and D_*m*_ are the diffusion constants for filaments and membrane in the cytosol, respectively. The drift velocities of filaments and membranes, designated by 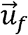 and 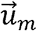, result from the pulling forces that are produced by molecular motors that link the filaments and the membranes:

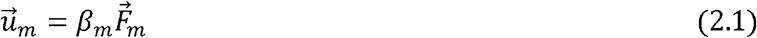

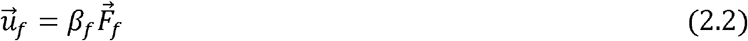

Where 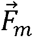 and 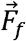 are the forces that act on the membrane and filaments, and *β_m_* and *β_f_* are the drag coefficients experienced by membrane fragments and filaments within the cytoplasm.

We assume that at low concentrations, no high-order structures are formed, and so we consider only pairwise interactions between filaments and membranes. Under such conditions, the range of the forces is limited to the average filament length, *L_f_*. The total force experienced by either the filaments or the membranes in a location 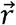 is then given by

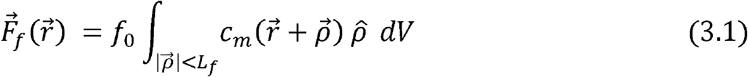

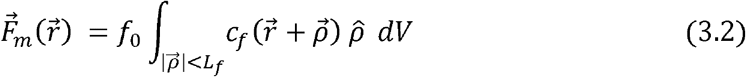

We assume cytoskeletal filaments polymerize at a fixed rate, *α*, such that

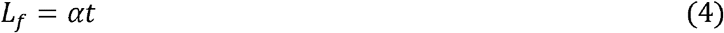

We substitute equations (2-4) into equations (1) and solve the resulting equations under boundary conditions of zero flux at the droplet surface, and initial conditions of homogeneous distribution. The solution is determined numerically by the finite-elements method, using the commercial software Comsol.

---

**Suppl. Fig. 1:** No cytoplasmic self-organization or centering in presence of latrunculin A and nocodazole which disrupt and prevent actin and microtubule filament formation, respectively. According to this data unlikely that e.g. intermediate filament systems involved.

**Suppl. Fig. 2:** Self-organization of cytoplasm in asymmetrically shaped microfluidic droplets. In dumbbell-shaped droplets, cytoplasmic self-organization robustly finds geometrical centroid. DiI for labeling Golgi and ER derived internal membranes (left panel), purified alexa647-tubulin (middle panel), and DIC images (right panel).

## References

1. Matthews, S., Observations on pigment migration within the fish melanophore. J Exp Zool, 1931. 58: p. 471–486.

2. McNiven, M.A., M. Wang, and K.R. Porter, Microtubule polarity and the direction of pigment transport reverse simultaneously in surgically severed melanophore arms. Cell, 1984. 37(3): p. 753–65.

3. McNiven, M.A. and K.R. Porter, Microtubule polarity confers direction to pigment transport in chromatophores. J Cell Biol, 1986. 103(4): p. 1547–55.

4. Rodionov, V.I. and G.G. Borisy, Self-centring activity of cytoplasm. Nature, 1997. 386(6621): p. 170–3.

5. Malikov, V., et al., Centering of a radial microtubule array by translocation along microtubules spontaneously nucleated in the cytoplasm. Nat Cell Biol, 2005. 7(12): p. 1213–8.

6. Borisy, G.G. and V.I. Rodionov, Lessons from the melanophore. FASEB J, 1999. 13 Suppl 2: p. S221–4.

7. Kimura, A. and S. Onami, Computer simulations and image processing reveal length-dependent pulling force as the primary mechanism for C. elegans male pronuclear migration. Dev Cell, 2005. 8(5): p. 765–75.

8. Kimura, K. and A. Kimura, Intracellular organelles mediate cytoplasmic pulling force for centrosome centration in the Caenorhabditis elegans early embryo. Proc Natl Acad Sci U S A, 2011. 108(1): p. 137–42.

9. Hamaguchi, M.S. and Y. Hiramoto, Analysis of the role of astral rays in pronuclear migration in sand dollar eggs by the colcemid-UV method Deveop Growth and Differ, 1986. 28: p. 143–156.

10. Chabin-Brion, K., et al., The Golgi complex is a microtubule-organizing organelle. Mol Biol Cell, 2001. 12(7): p. 2047–60.

11. Rivero, S., et al., Microtubule nucleation at the cis-side of the Golgi apparatus requires AKAP450 and GM130. EMBO J, 2009. 28(8): p. 1016–28.

12. Efimov, A., et al., Asymmetric CLASP-dependent nucleation of noncentrosomal microtubules at the trans-Golgi network. Dev Cell, 2007. 12(6): p. 917–30.

13. Miller, P.M., et al., Golgi-derived CLASP-dependent microtubules control Golgi organization and polarized trafficking in motile cells. Nat Cell Biol, 2009. 11(9): p. 1069–80.

14. Vinogradova, T., et al., Concerted effort of centrosomal and Golgi-derived microtubules is required for proper Golgi complex assembly but not for maintenance. Mol Biol Cell, 2012. 23(5): p. 820–33.

15. Rodionov, V.I., et al., Functional coordination of microtubule-based and actin-based motility in melanophores. Curr Biol, 1998. 8(3): p. 165–8.

16. Schuh, M., An actin-dependent mechanism for long-range vesicle transport. Nat Cell Biol, 2011. 13(12): p. 1431–6.

17. Waterman-Storer, C., et al., Microtubules remodel actomyosin networks in Xenopus egg extracts via two mechanisms of F-actin transport. J Cell Biol, 2000. 150(2): p. 361–76.

18. Holtze, C., et al., Biocompatible surfactants for water-in-fluorocarbon emulsions. Lab Chip, 2008. 8(10): p. 1632–9.

19. Pinot, M., et al., Effects of confinement on the self-organization of microtubules and motors. Curr Biol, 2009. 19(11): p. 954–60.

20. Anna, S.L., N. Bontoux, and H.A. Stone, Formation of dispersions using “flow focusing” in microchannels. Applied Physics Letters, 2002. 82: p. 364–367.

21. Garstecki, P., H.A. Stone, and G.M. Whitesides, Mechanism for flow-rate controlled breakup in confined geometries: a route to monodisperse emulsions. Phys Rev Lett, 2005. 94(16): p. 164501.

22. Jimenez, A.M., et al., Towards high throughput production of artificial egg oocytes using microfluidics. Lab Chip, 2011. 11(3): p. 429–34.

23. Kirschner, M.W. and T. Mitchison, Microtubule dynamics. Nature, 1986. 324(6098): p. 621.

24. Lenart, P., et al., A contractile nuclear actin network drives chromosome congression in oocytes. Nature, 2005. 436(7052): p. 812–8.

25. Joanny, J.F. and J. Prost, Active gels as a description of the actin-myosin cytoskeleton. HFSP J, 2009. 3(2): p. 94–104.

26. Tee, Y.H., et al., Cellular chirality arising from the self-organization of the actin cytoskeleton. Nat Cell Biol, 2015. 17(4): p. 445–57.

27. Desai, A., et al., The use of Xenopus egg extracts to study mitotic spindle assembly and function in vitro. Methods Cell Biol, 1999. 61: p. 385–412.

28. Burkel, B.M., G. von Dassow, and W.M. Bement, Versatile fluorescent probes for actin filaments based on the actin-binding domain of utrophin. Cell Motil Cytoskeleton, 2007. 64(11): p. 822–32.

29. Field, C.M., et al., Actin behavior in bulk cytoplasm is cell cycle regulated in early vertebrate embryos. J Cell Sci, 2011. 124(Pt 12): p. 2086–95.

30. Garstecki, P., et al., Formation of monodisperse bubbles in a microfluidic flow-focusing device. Applied Physics Letters, 2004. 85(13): p. 2649–2652.

