## Supplementary figures and images for "Cytoplasmic self-organization established by internal lipid membranes in the interplay with either actin or microtubules"

### Supplemental Figure 1

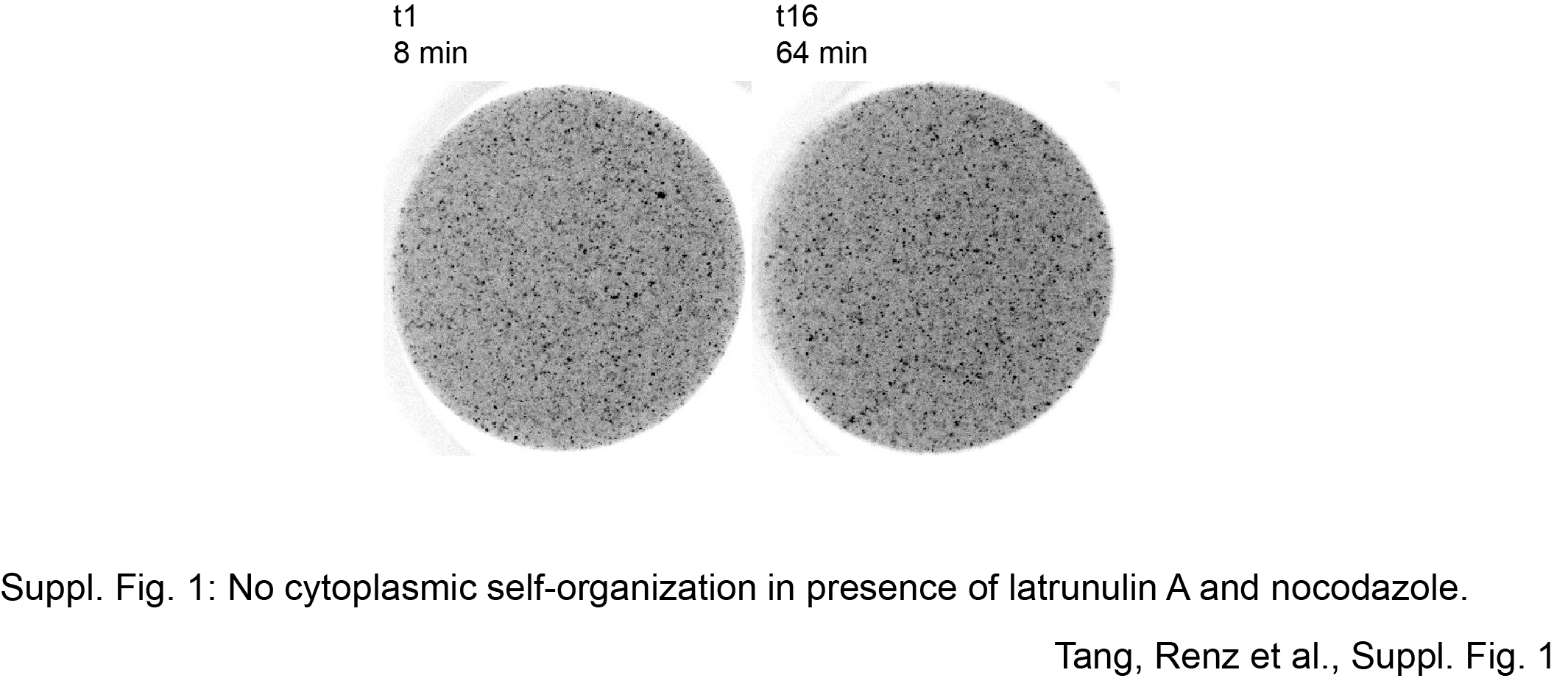

### Supplemental Figure 2

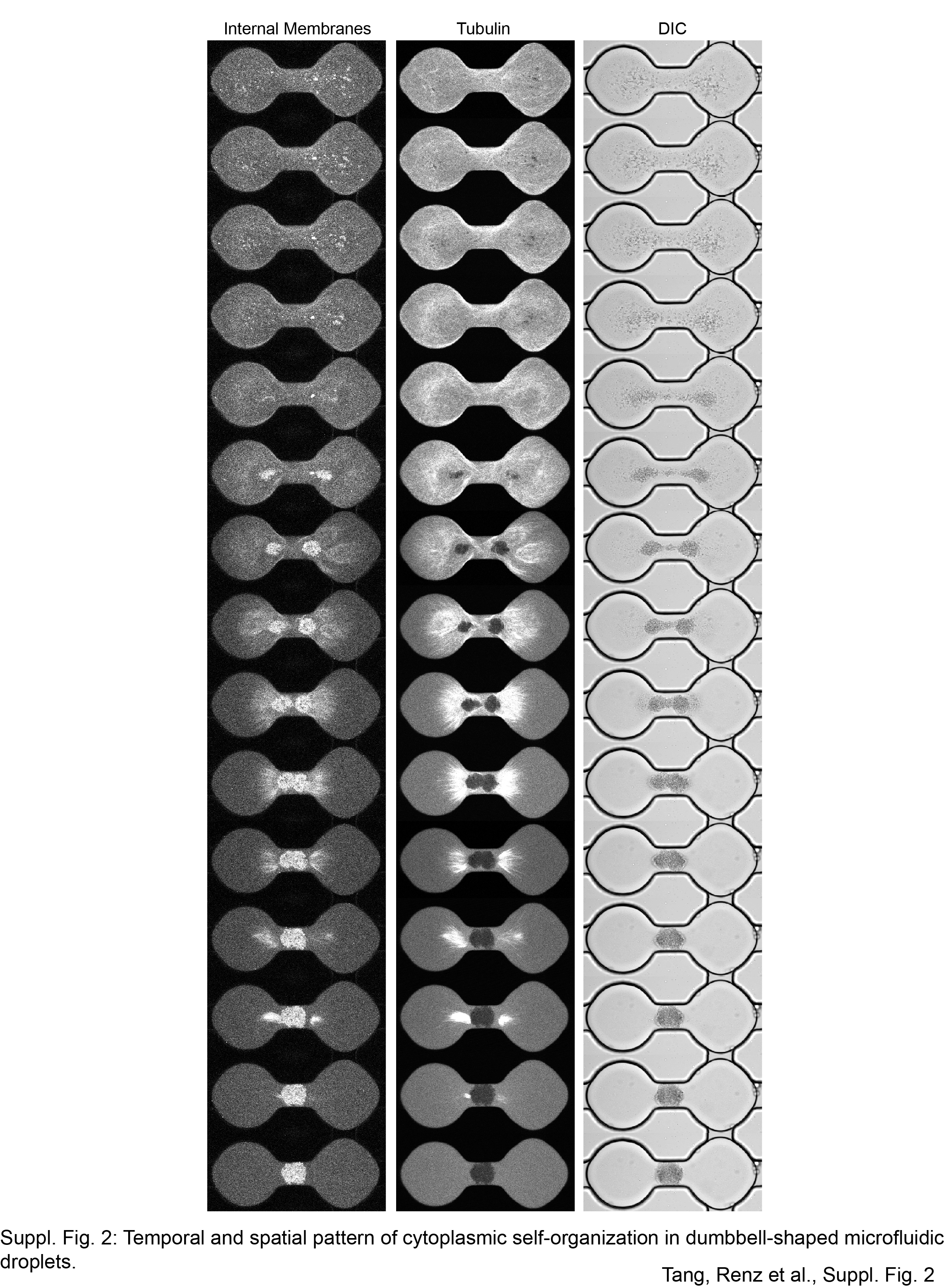
